# Social Anxiety Increases Autonomic and Visuocortical Generalization of Conditioned Aversive Responses to Faces

**DOI:** 10.1101/2025.07.14.663966

**Authors:** Jourdan J. Pouliot, Richard T. Ward, Faith Gilbert, Payton Chiasson, Caitlin Traiser, Andreas Keil

## Abstract

Aversive generalization learning is an adaptive trait that is necessary for survival in a dynamic environment. However, this process is exaggerated in persons with anxiety disorders, leading to overgeneralization of learned threat associations, hyperreactive fight-or-flight responses, and persistent avoidance. Patients with social anxiety disorder (SAD) exhibit impaired conditioned threat discrimination particularly with respect to social stimuli, such as faces. The present study examined the relationship between social anxiety and generalization of visuocortical and pupil dilation responses to a series of facial morphs, one of which was always paired with a noxious sound. Steady-state visual evoked potentials (ssVEPs; N = 65) increasingly fit a model of generalization, and pupil dilation responses (N = 62) also decreasingly discriminated the CS+ as a function of social anxiety. These results contribute to a growing body of work suggesting that SAD dysregulates the ability of autonomic responses to specifically target social threat. The finding of widened visuocortical tuning in SAD implicates a role of the visual system in driving attentional biases in anxiety disorders, including increased visual processing of safety signals similar to threat cues.

## Introduction

The ability to form aversive memories in response to threats and generalize to similar situations is an adaptive trait necessary for navigating the world (LeDoux, 2003). However, dysregulation of processes linked to fear memory acquisition and fear generalization has been theorized to underlie the etiology of anxiety disorders (Mineka & Zinbarg, 2006; Fullana et al., 2020). These disorders are characterized by excessive fears of certain stimuli or situations accompanied by avoidance, heightened arousal, and distress (Hoehn-Saric & McLeod, 2000; Szuhany & Simon, 2022). Additionally, anxiety disorders have been shown to impair working memory and one’s capacity to flexibly allocate attention, which are necessary for successfully navigating dynamic environments (Robinson et al., 2013; Eysenck et al., 2007).

Pavlovian conditioning has been identified as the central mechanism underlying the etiology and maintenance of anxiety disorders (Pavlov, 1927; Lissek et al., 2005). It is characterized as the formation of representations comprised of logical and perceptual associations between stimuli and situations which are informed by preconceptions (Rescorla, 1988). An extension of Pavlovian conditioning is stimulus generalization learning, which extends associations between a conditioned stimulus (CS+) and outcome (unconditioned stimulus, US) across a gradient of similar stimuli. Aversive generalization— involving an unpleasant US—facilitates graded defensive responses following exposure to stimuli along a given dimension defined by similarity to the CS+. These generalization stimuli (GS) may trigger conditioned fear responses despite never preceding the aversive outcome.

Aversive generalization has been found to be exaggerated in individuals with anxiety disorders, specific phobia, and post-traumatic stress disorder, and is thought to reflect impaired discrimination between threat and safety cues (Lenaert et al., 2014; Dymond et al., 2015; Jovanovic et al., 2012). Accordingly, autonomic responses such as heart rate and the startle-blink reflex exhibit greater generalization of conditioned fear responses to unpaired stimuli in socially anxious people compared to healthy controls (Lissek et al., 2010; Ahrens et al., 2016). Studies measuring conditioned pupillary responses have found increased pupil dilation in response to the CS+, but mixed results supporting a generalization response gradient and the effects of anxiety disorders on such a gradient (Leuchs et al., 2017; Reutter & Gamer, 2023; Reutter et al., 2025; Greenberg et al., 2013; Farkas et al., 2024).

Neuroscience studies have found that sensory cortices contribute to the memorization and expression of conditioned fear (Miskovic & Keil, 2012; Dolan, Heinze, Hurlemann, & Hinrichs, 2006; Bröckelmann et al., 2011; Weinberger, 2011). Work utilizing electroencephalography (EEG) has found enhanced power in the steady-state visual potential (ssVEP) response to the CS+, compared to the CS– (Miskovic & Keil, 2013; Wieser et al., 2014; Kastner et al., 2015; Petro et al., 2017). Aversive generalization studies using Gabor patches found, in occipital regions, the greatest ssVEP activity in response to the CS+ and suppressed ssVEP activity to GS that were most similar in orientation to the CS+(McTeague et al., 2015; Antov et al., 2020; Friedl & Keil, 2021). This sharpening pattern (also known as a “Mexican Hat” pattern) has been linked to lateral inhibitory interactions between orientation sensitive cells in V1 neurons, although little research has been done to identify this mechanism at the neural population level (Blakemore & Tobin, 1972; Freeman et al., 2002; Shapley et al., 2007). Alongside this finding, however, studies have found generalization of the ssVEP signal across the stimulus gradient in parietal channels (McTeague et al., 2015).

Few aversive generalization studies utilizing ssVEPs have used stimuli with greater semantic content, such as faces. Unlike simple stimuli, perception of faces involves an integration of disparate facial parts and has been localized to right occipitotemporal channels (Ales et al., 2012; Boremanse et al., 2013; Boremanse et al., 2014). One study presenting facial stimuli varying in emotional expression (angry, fearful, happy, neutral) to treatment-seeking patients and control participants found that angry and fearful expressions led to an increased parieto-occipital response, relative to neutral faces (McTeague et al., 2018). Notably, this effect was greatest in patients with social anxiety disorder (SAD), relative to patients with other anxiety disorders and healthy controls. A subsequent study employed an aversive generalization paradigm, wherein both facial identity and emotional expression were varied and subjects varied in social anxiety intensity (Stegmann et al., 2020). This study found a sharpening pattern in low-level visual cortex that was enhanced as social anxiety increased. Altogether, this suggests that social anxiety amplifies neural responses to emotional faces and increases suppression of similar faces during conditioned threat paradigms.

The present study is a replication and extension of Stegmann et al. (2020) with changes made to facilitate response generalization as opposed to sharpening. First, Stegmann et al. (2020) used a dedicated acquisition phase, wherein participants learned to discriminate the CS+ from a CS– when presented alone. The present study did not include a separate acquisition phase but instead began with generalization conditioning, with the GS and CS+ faces shown intermittently, and only the CS+ face paired. Such an approach has been shown to diminish categorical perception of the CS+ and hence reduce discrimination, heightening generalization (Asok et al., 2019; Dunsmoor & Murphy, 2015). Additionally, participants in the present study were only presented with emotionally neutral facial stimuli, whereas the CS+ face used by Stegmann et al. (2020) presented a fearful expression during presentation of the US. Lastly, participants were conditioned to a bilateral gradient of emotionally neutral faces, across a large number of trials, also thought to facilitate generalization in the animal model (Asok et al., 2019).

This study employed multiple behavioral and physiological correlates to investigate the impact of social anxiety on conditioned fear generalization, including self-report measures, pupil dilation, and ssVEP. To assess patterns of generalization learning, the inner product was taken between observed behavioral and physiological response gradients and hypothesized weight vectors. In keeping with previous findings, it was hypothesized that greater social anxiety would increase the sharpening of visuocortical ssVEP response patterns. For comparison, and based on changes made to the experimental design, increased generalization of ssVEP responses as a function of social anxiety was also examined as a competing hypothesis. In line with autonomic evidence, it was hypothesized that social anxiety would also lead to broader generalization patterns of pupil dilation.

Social anxiety was also hypothesized to be positively related to broader generalization patterns of self-reported valence and arousal across the stimulus gradient. To test the competing hypothesis that motor responses increasingly discriminate the CS+ as social anxiety increases, pupillary responses and self-report data were also fit to an all-or-nothing model. This hypothesis was informed by previous findings that pupil responses and self-report ratings discriminated the CS+, rather than generalize across the stimulus gradient (Farkas et al., 2024). Because there is no known mechanism for evoking a sharpening pattern in behavioral and pupillary response functions, this model was not fit to pupil responses or self-report ratings.

## Methods

### Participants

The study was pre-registered at OSF (https://osf.io/quy47). The present manuscript reports on a continuous analysis of social anxiety, pre-registered as an exploratory analysis. We determined a required sample size of 62 participants by Monte-Carlo simulations (Boudewyn et al., 2018; Gibney et al., 2020). A total of 83 participants (68 female) from the University of Florida and members of the community participated in this study either for course credit or $70 monetary compensation. Advertisements targeted individuals seeking treatment for social anxiety, and recruitment also included the specialty clinics of the University of Florida. Participants were at least 18 years old and were screened for uncorrected visual impairments and epilepsy or family history of epilepsy. 12 participants withdrew from the study due to overwhelming discomfort with the US. 1 participant did not complete the questionnaires and so was excluded from all analyses. Behavioral data from the remaining 70 participants (55 females, M_AGE_ = 21.14, SD_AGE_ = 5.16) were used for analyses. Of the participants included in behavioral analyses, 54 identified as White (77.14%), 5 identified as Black (7.14%), 6 identified as Asian (8.57%), 5 identified as Mixed Race/Other (7.14%). For the EEG analyses, 5 participants used for the behavioral analyses were excluded from EEG data analyses due to excessive noise in EEG data, leaving 65 participants (50 females, M_AGE_ = 20.80, SD_AGE_ = 4.51) remaining. Of those included in the EEG analyses, 50 participants identified as White (76.92%), 4 identified as Black (6.15%), 6 identified as Asian (9.23%), 5 identified as Mixed Race/Other (7.69%). For the pupillary response analyses, 8 participants used for the behavioral analyses were excluded from the pupillary response analyses due to excessive noise in the pupil data, leaving 62 participants (50 females, M_AGE_ = 20.80, SD_AGE_ = 4.51) remaining. Of those included in the pupillary response analyses, 50 participants identified as White (81.65%), 4 identified as Black (6.45%), 3 identified as Asian (4.84%), 5 identified as Mixed Race/Other (8.06%). This study was approved by the local Institutional Review Board and was conducted in accordance with the Declaration of Helsinki.

All participants completed the Liebowitz Social Anxiety Scale (LSAS) prior to the experimental session as part of a screening procedure intended to increase variability in the covariate of interest (Liebowitz, 1987). Effort was taken to recruit participants scoring high (>80) in the LSAS total score. Participants completed the LSAS questionnaire again during the session prior to completing the experimental paradigm. This latter score was used as a predictor in all continuous analyses described below. Participants also completed additional surveys including the Beck Depression Inventory (BDI), the Mood and Anxiety Symptom Questionnaire (MASQ), the State-Trait Anxiety Inventory (STAI), the Posttraumatic Diagnostic Scale (PDS-5), and the EASI Temperament Survey, although these were not incorporated into the analyses.

### Materials and Stimuli

Visual stimuli were presented on a Display++ monitor (Cambridge Research Systems Ltd.) at a refresh rate of 120Hz. Conditioned stimuli consisted of seven monochrome female faces (Figure 1). A Gaussian filter (FWHM = 50 grayscale units) was applied to all faces to correct for differences in contrast. Three of these faces (GS1, CS+, GS6) were drawn from the Karolinska Directed Emotional Faces (KDEF) database. The other facial stimuli were generated by systematically morphing the original faces using the MorphX software (https://www.norrkross.com/). In particular, the GS2 and GS3 faces were, respectively, 70:30 and 30:70 morphs of the GS1 and CS+. Likewise, the GS4 and GS5 were, respectively, 70:30 and 30:70 morphs of the CS+ and GS6. All facial stimuli were flickered at a driving frequency of 15Hz to elicit an ssVEP response. Self-Assessment Manikins (SAMs) were presented at certain points throughout the experiment (Bradley & Lang, 1994). SAMs are 9-point scales used to assess self-reported valence and arousal in response to viewing each facial stimulus. The unconditioned stimulus (US) was a burst of 92dB(A) white noise. The noise was played bilaterally from speakers placed behind the participant.

**Figure 1.**
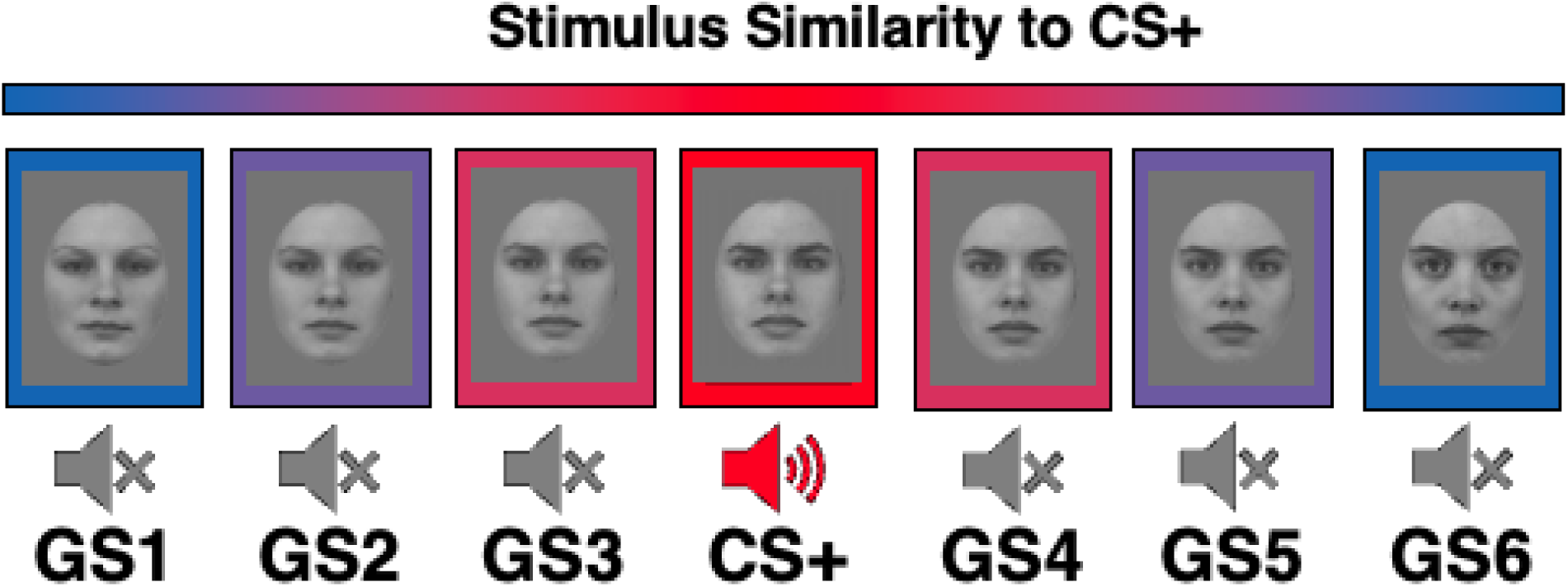
*Stimulus Gradient*. Three facial stimuli (GS1, CS+, GS6) were sourced from the Karolinska Directed Emotional Faces (KDEF) database. The remaining stimuli were derived from 30-70 morphs of the original faces.

### Design and Procedure

During the experiment, participants were seated approximately 120cm from the display monitor and 60cm from the eye tracker. The paradigm was programmed using the MATLAB Psychophysics Toolbox (Brainard, 1997; Pelli, 1997). The paradigm contained 291 trials. During each trial, one of the conditioned stimuli was presented by the display monitor at a visual angle of approximately 7.63°. The first 11 trials were booster trials which presented either the GS1, CS+, or GS6 to elicit familiarity with the endpoints of the gradient and thereby facilitate generalization learning. The remaining 280 trials sampled each of the seven facial stimuli pseudorandomly such that each stimulus was presented 40 times.

Following an intertrial interval (ITI) ranging from 2-4s seconds, each GS was presented for 2s (Figure 2, left). The CS+ was presented for 3s and co-terminated with presentation of the US, which played during the last second (Figure 2, right). SAMs were presented at a visual angle of approximately 21.47° beneath each facial stimulus at three different points in the experiment: prior to the start of conditioning to collect baseline ratings, following an early trial (trial 87), and following a later trial (trial 168).

**Figure 2.**
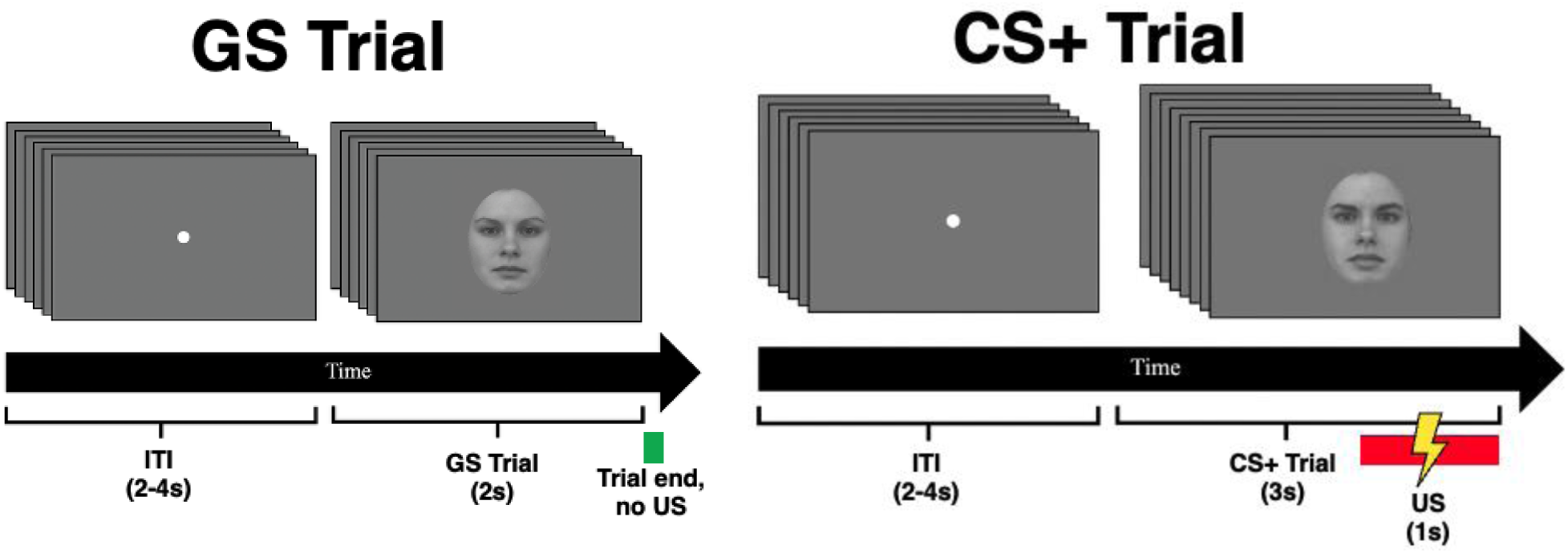
*Trial Time Course*. During a CS+ trial (*Right*), the CS+ was presented for 3 seconds, co-terminating with a noxious sound (unconditioned stimulus, US, 1s duration). Only the first 2 seconds were used for analysis. During any other trial type (*Left*), a generalization stimulus (GS) was presented for 2 seconds and no US occurred. Both trial types were preceded by an inter-trial interval with a random duration from 2 to 4 seconds.

### Data Collection and Processing

EEG was collected using a 128-channel system from Electrical Geodesics, Inc. (EGI). During data collection, electrical signals were recorded from the scalp at a sampling rate of 500Hz using electrode Cz as an online reference. Prior to the start of the experiment, electrode impedances were kept below 60kΩ.

EEG data were preprocessed in MATLAB using the open-source Electro/MagnetoEncephalography Software (EMEGS; Peyk, De Cesari, & Junghofer, 2011). Trials were segmented such that segments began 800ms pre-stimulus and extended until 2000ms post-stimulus. A 40Hz (3dB point) low-pass Butterworth filter (45 dB/octave, 23^rd^ order) and a 4Hz (3 dB point) high-pass Butterworth filter (18 dB/octave, 2^nd^ order) were applied to each segment. Signals were re-referenced offline to the average signal across all channels. Then an artifact rejection algorithm was applied to the data (Junghöfer, Elbert, Tucker, & Rockstroh, 2000). Artifactual data were identified in two steps based on the scalp-wide distribution of voltage characteristics, namely the absolute mean, the standard deviation, and gradient. First, global bad channels were identified as being greater than 3 standard deviations above the median of the distributions previously mentioned, across all trials. These channels were removed and replaced with spherical spline-interpolated values. Second, bad channels in each trial were identified as being greater than 3 standard deviations above the median of the aforementioned distributions, within each trial. A maximum number of 20 channels was interpolated for each trial, after which the trial was discarded. The distribution of bad channels was manually inspected for each trial to check that they were not disproportionately localized in a particular cluster of electrodes. Eye movement artifacts were subsequently removed using a regression-based algorithm (Schlögl et al., 2007). This approach identified eye movements from the HEOG and VEOG channels before subtracting them from the scalp-wide EEG data. Lastly, to reduce the effects of volume conduction and thereby improve localization of signals from the surface of the brain, the current source density (CSD) transformation was applied to the preprocessed EEG data (Junghöfer et al., 1997).

Pupil data were collected using an Eyelink 1000 Plus SR Research camera with a 16mm lens. During recording, the camera was set to “ellipse mode” and illumination of the infrared signal was set to 100%. Pupil data were processed using custom MATLAB scripts. Data for each trial were segmented such that each segmented began 1000ms prior to stimulus onset and extended until 2000ms post-stimulus onset. A pupil artifact rejection algorithm identified segments as artifactual if pupil diameter ever reached zero, if the pupil diameter change across two sample points exceeded 2.5mm, or if low-frequency drift across two sample points (extracted using a 7.5Hz (6^th^ order) low-pass filter) exceeded 5mm.

### Data Reduction and Analysis

Prior to statistical analysis, psychophysiological signals were further processed and averaged. With respect to the EEG data, following condition-wise averaging of trials and baseline subtraction (baseline period: –438 to –2ms), the Hilbert transform (9^th^ order Butterworth) was used to extract the time-varying 15Hz amplitude envelope (Figure 3).

**Figure 3.**
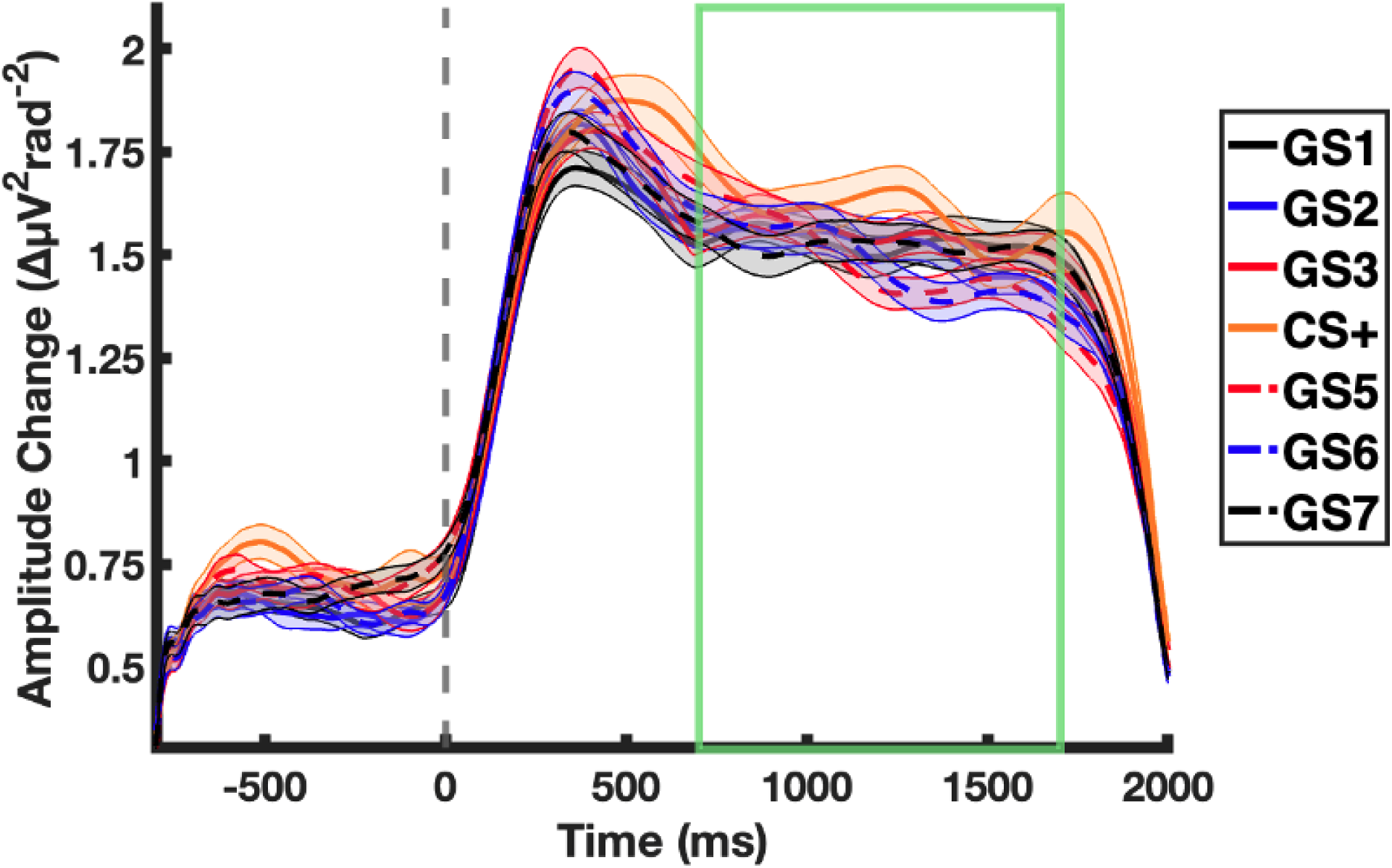
*Grand mean (n=65) ssVEP Hilbert Transform Time Series*. All facial stimuli were flickered at 15Hz to elicit a steady-state visual evoked potential (ssVEP). To isolate this waveform from condition averages, time-averaged baseline data (baseline period: –360ms to –2ms) were subtracted from the time series and a Hilbert Transform (9^th^ order Butterworth) was applied to the data. Shown here are the condition-wise average results at channel POz. For all ssVEP analyses, Hilbert data from each condition were time-averaged from +700ms to +1700ms (*green window*). Shaded error bars display within-subject variability (Cousineau, 2005).

Time-frequency data were averaged from +700ms to +1700ms, yielding, for each participant, a gradient of average ssVEP amplitudes across facial stimuli. Condition averaged pupil size data were also subjected to baseline subtraction (baseline period: – 360ms to –100ms), prior to averaging across the period from +1500ms to +2000ms (Figure 4).

**Figure 4.**
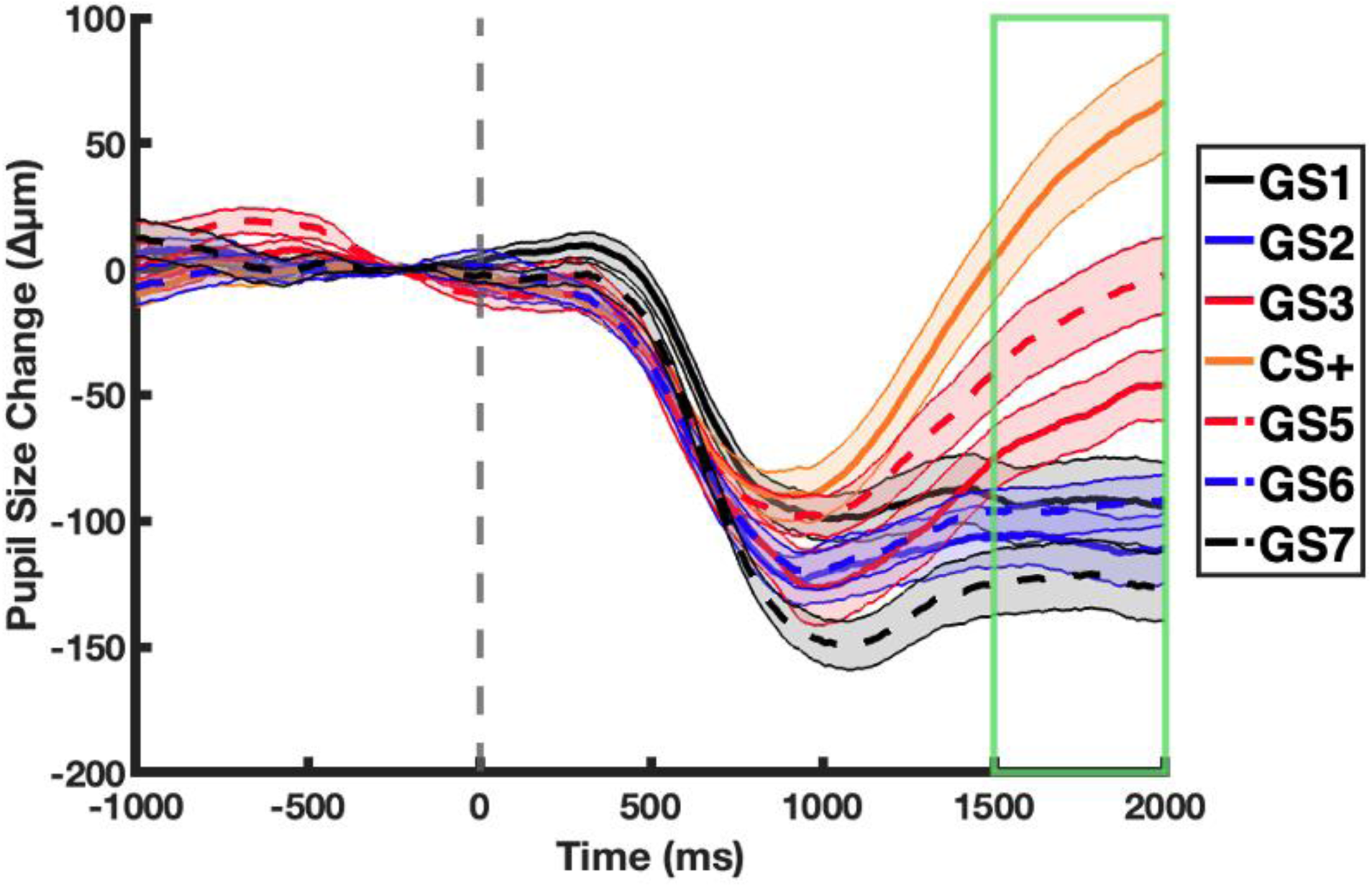
*Grand mean (n=62) Pupil Size Time Series.* Following pre-processing, time-averaged baseline data (baseline period: –360ms to –100ms) were subtracted from condition averaged pupil size data. For all pupil dilation analyses, pupil size data from each condition were time-averaged from +1500ms to +2000ms (*green window*). Shaded error bars display within-subject variability (Cousineau, 2005).

Valence and arousal ratings, averaged pupil responses, and averaged ssVEP responses were fit to the a priori models and correlated with LSAS ratings. Adherence of observed responses across stimuli to one of the a priori models was assessed by computing the inner product between the response gradient and the corresponding weight vector (Figure 5). The ssVEP response gradient for each channel and each participant was multiplied, as an inner product, with the generalization weight vector ([–3.0, 0.5, 1.5, 2.0, 1.5, 0.5 –3.0]) and the sharpening weight vector ([0.5, –1.0, –2.0, 5.0, –2.0, –1.0, 0.5]) to yield model fit metrics for these models. The pupil response gradient for each participant was multiplied by the generalization weight vector and the all-or-nothing weight vector ([– 1.0, –1.0, –1.0, 6.0, –1.0, –1.0, –1.0]). The early and late valence and arousal ratings for each participant were also multiplied by the generalization weight vector and all-or-nothing weight vector.

**Figure 5.**
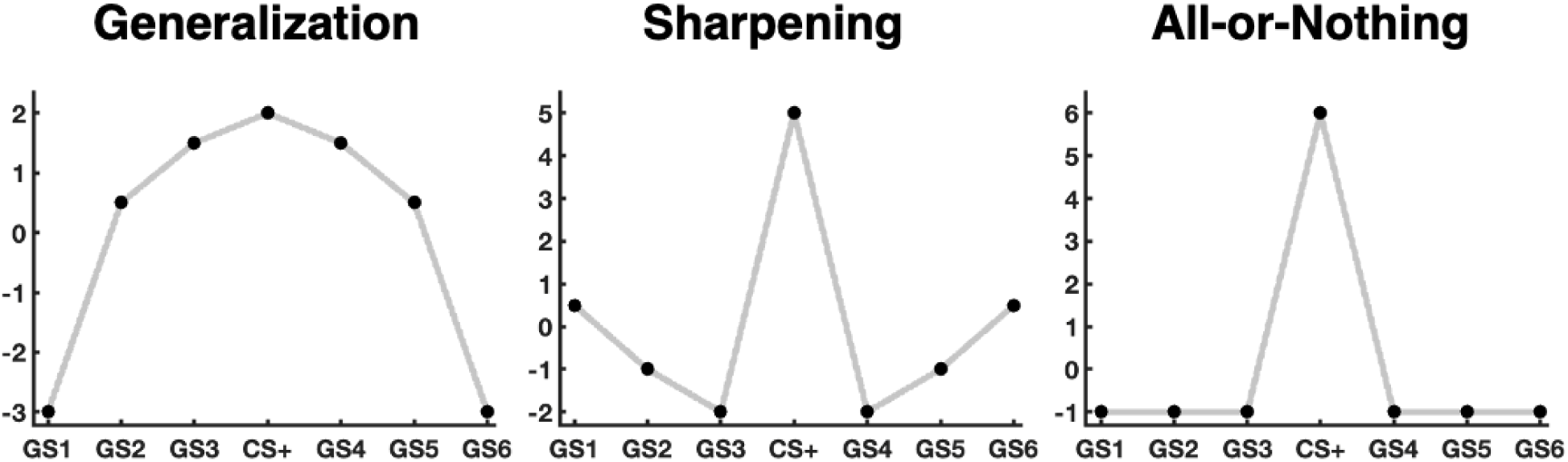
*A Priori Models*. The shapes of the observed psychophysiological and behavioral response patterns were characterized by taking the inner product of each participant’s condition-wise average responses and two of the above weight vectors. The ssVEP response patterns were multiplied with the generalization model (*Left*) and the sharpening model (*Middle*). All other variables (pupil, valence, arousal) were multiplied with the generalization model (*Left*) and the all-or-nothing model (*Right*).

The inner products corresponding to each measure and model were then Z-scored across participants and correlated with LSAS scores. To account for the possibility of a non-linear but monotonic trend, Spearman’s ρ was used to measure the relationship between the ranks of the inner products and LSAS scores. To correct for multiple comparisons, permutation tests were conducted to determine critical ρ thresholds (Karniski, Blair, & Snider, 1994). Two-tailed tests were run to the increase robustness of hypothesized findings and consider results contrary to our hypotheses. Two critical ρ’s (2.5 and 97.5 percentiles of the permutation distribution) were obtained from each correlation analysis to determine significance.

Additional follow-up analyses were performed on inner products that were significantly correlated with social anxiety. Inner products were grouped into quintiles based on the corresponding LSAS scores. A linear F-contrast was then fit to the quintiles. In line with our hypotheses, it was predicted that increases in LSAS would lead to linear increases in the hypothesized model fit.

## Results

Averaged ssVEP responses for the GS1, CS+, and GS6 conditions (shown in Figure 6) localize most of the signal to a medial occipitoparietal cluster. ssVEP responses were expected to exhibit increased sharpening or generalization as a function of LSAS ratings (M = 54.57, SD = 23.89, Min = 7, Max = 105). Results did not show a significant positive relationship between LSAS scores and fit to the sharpening model (all p’s > 0.025). Instead, a significant positive correlation was found between LSAS scores and fit to the generalization model. This was found for a right occipitoparietal cluster of 3 channels (ρ = 0.38, ρ = 0.37, ρ = 0.40; all p’s < 0.025) (Figure 7). As a follow-up analysis, response gradients were averaged within this cluster, and the inner products between the cluster-averaged ssVEP response gradients and the generalization model were split into quintiles based on LSAS score (Figure 8, Supplementary Figure 1). A significant linear trend was found across these quintiles (F(4, 61) = 9.28, p < 0.00001) (Figure 9). This indicates that increased social anxiety was associated with increased generalization of the ssVEP response gradient in right occipitoparietal channels.

**Figure 6.**
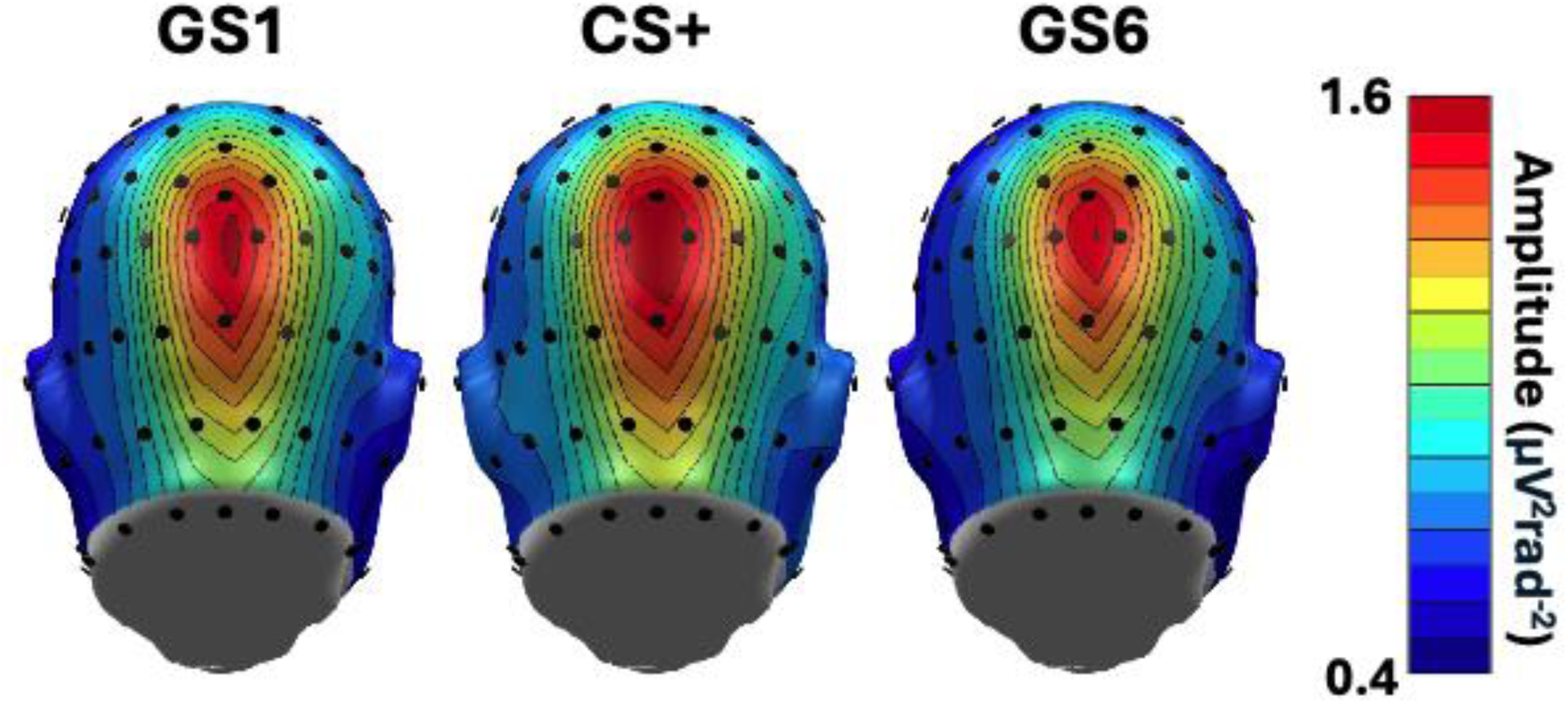
*Grand mean ssVEP amplitude Topographies*. The topographies show pronounced amplitude maxima at medial occipitoparietal channels. In these channels, responses were greatest following presentation of the CS+.

**Figure 7.**
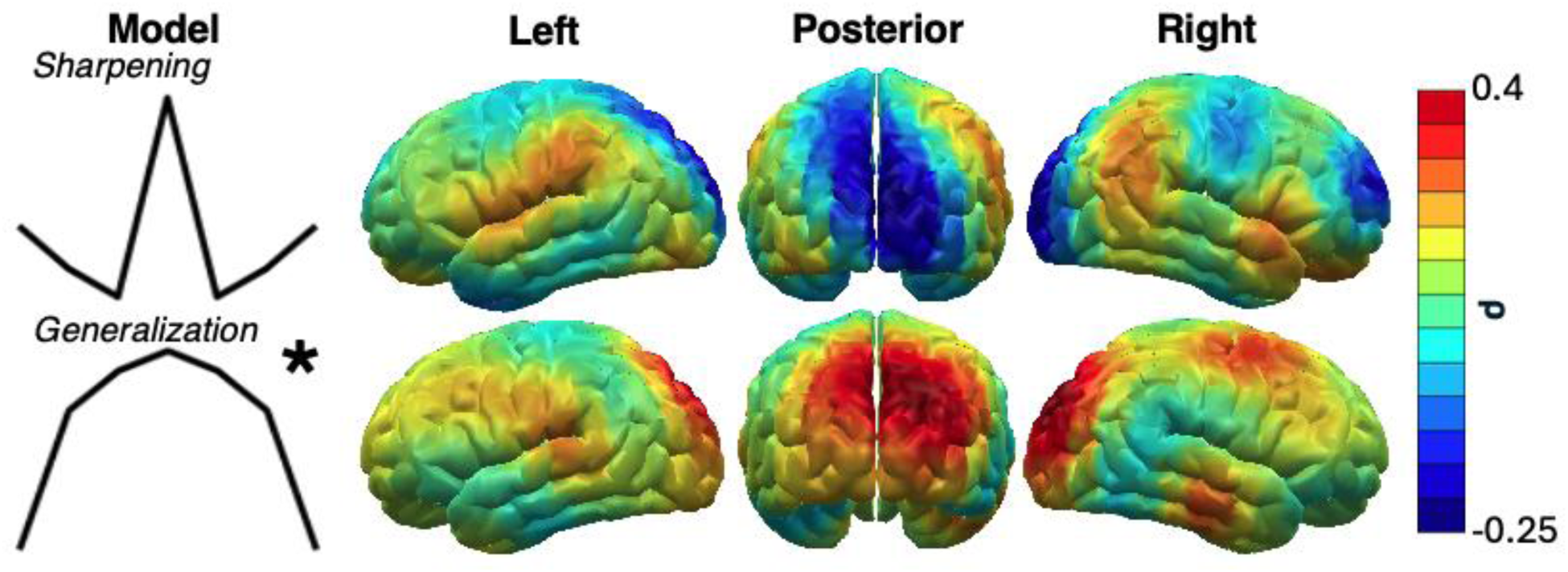
*Rank-Correlation Topographies Between the ssVEP Model Fit (in CSD space) and Social Anxiety Ratings*. Mass-univariate rank-correlations found that social anxiety was significantly correlated with a cluster of medial and right-of-center channels. * indicates p < 0.025.

**Figure 8.**
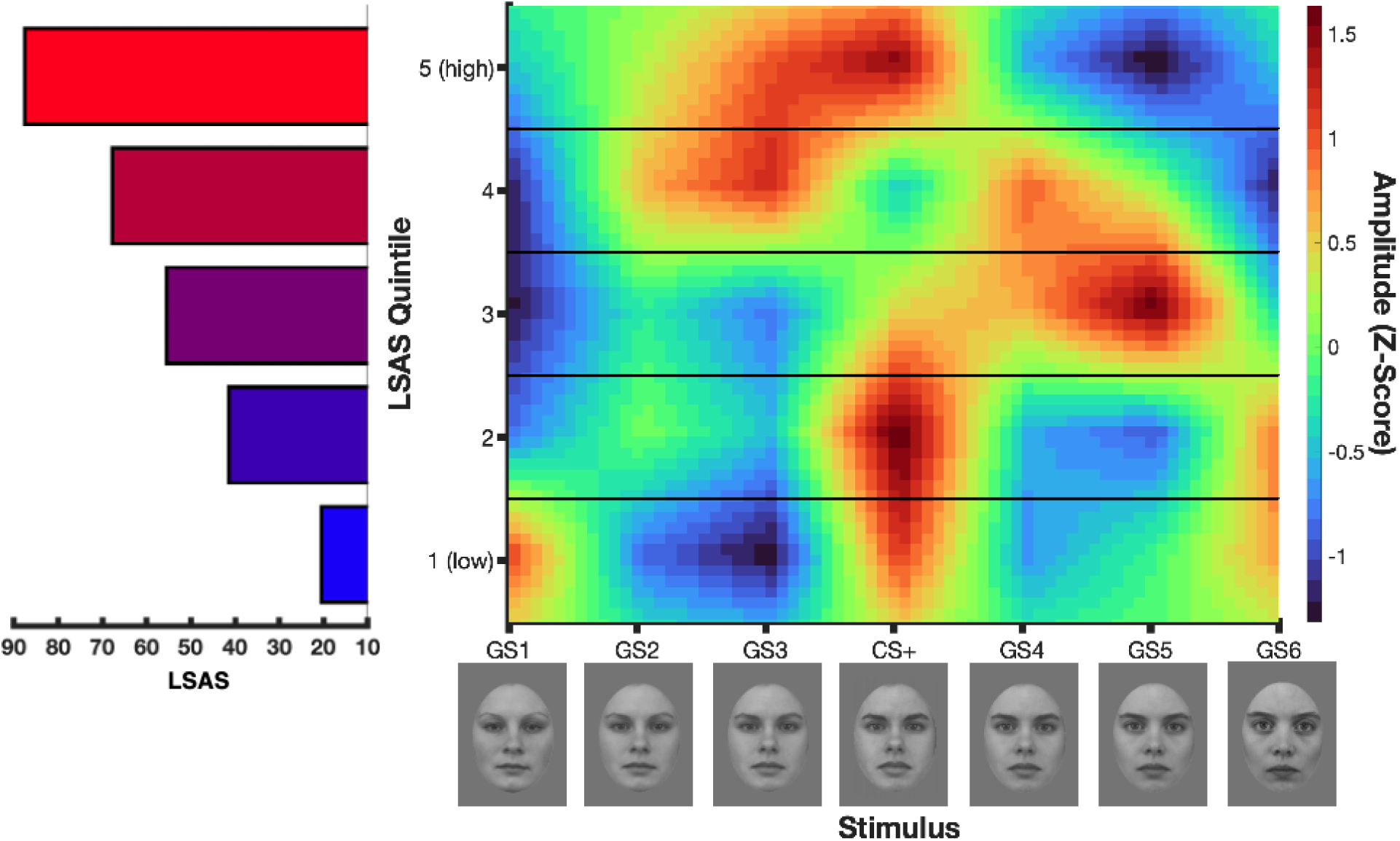
*ssVEP Response Gradients Grouped by LSAS Quintile*. ssVEP data were separated into quintiles with respect to LSAS ratings (*Left*). Each quintile average response gradient was Z-scored to enable comparison between groups (*Right*).

**Figure 9.**
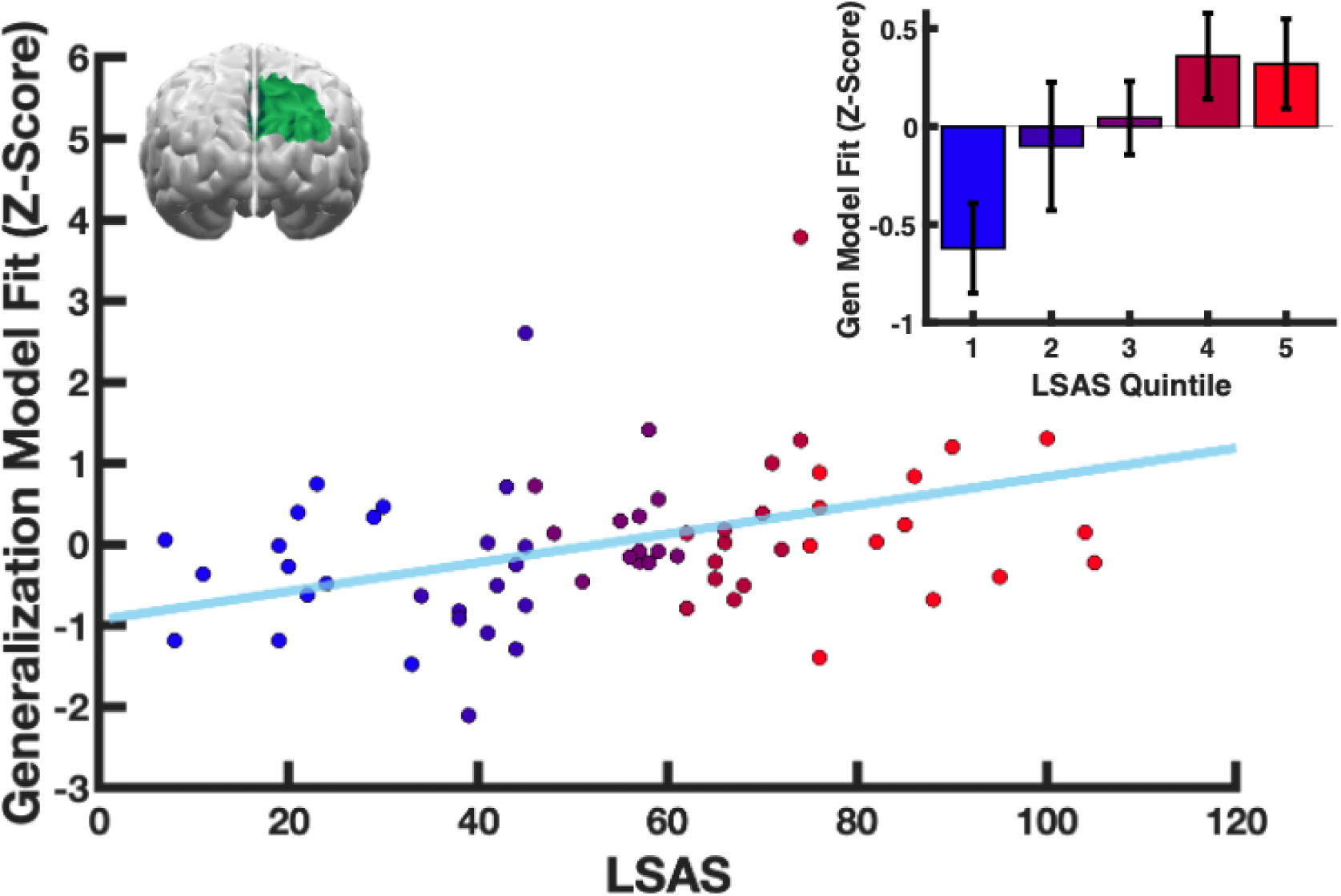
*Occipitoparietal Response Gradient Fit to Generalization Model Against LSAS Ratings*. Visuocortical ssVEP response patterns from channels exhibiting significant rank-correlations with social anxiety ratings were averaged across channels (right occipitoparietal channels, *Top Left*). The inner product was taken between each participant’s condition-wise response pattern and the generalization pattern. The inner products were then Z-scored across participants, plotted against LSAS ratings, and color-coded based on LSAS quintile. Z-scored inner product averages and standard errors were calculated with respect to each LSAS quintile (*Top Right*).

Pupillary responses were expected to exhibit increased generalization or discrimination as a function of LSAS ratings (M = 53.76, SD = 23.50, Min = 7, Max = 105). While the former was not shown to be the case (ρ = –0.20, p > 0.025), a significant negative correlation was found between LSAS scores and fit to the all-or-nothing model (ρ = –0.31, p < 0.025). The inner products between the pupil response gradient and the all-or-nothing model were split into quintiles based on LSAS score (Figure 10, Supplementary Figure 2). A significant linear trend was found across quintiles (F(4, 57) = 6.20, p < 0.001; Figure 11).

**Figure 10.**
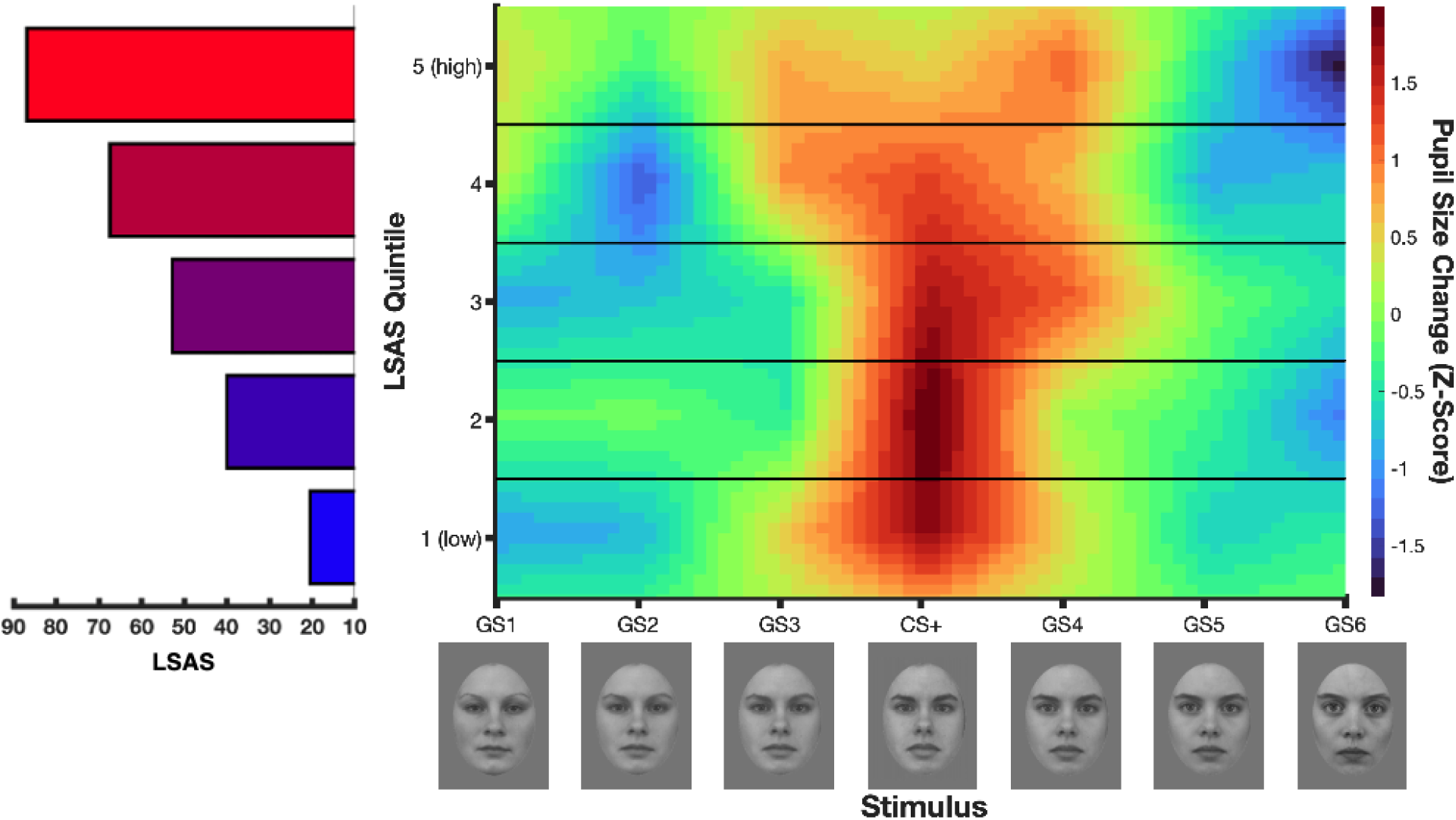
*Pupil Dilation Response Gradients Grouped by LSAS Quintile.* Pupil data were separated into quintiles with respect to LSAS ratings (*Left*). Each quintile average response gradient was Z-scored to enable comparison between groups (*Right*).

**Figure 11.**
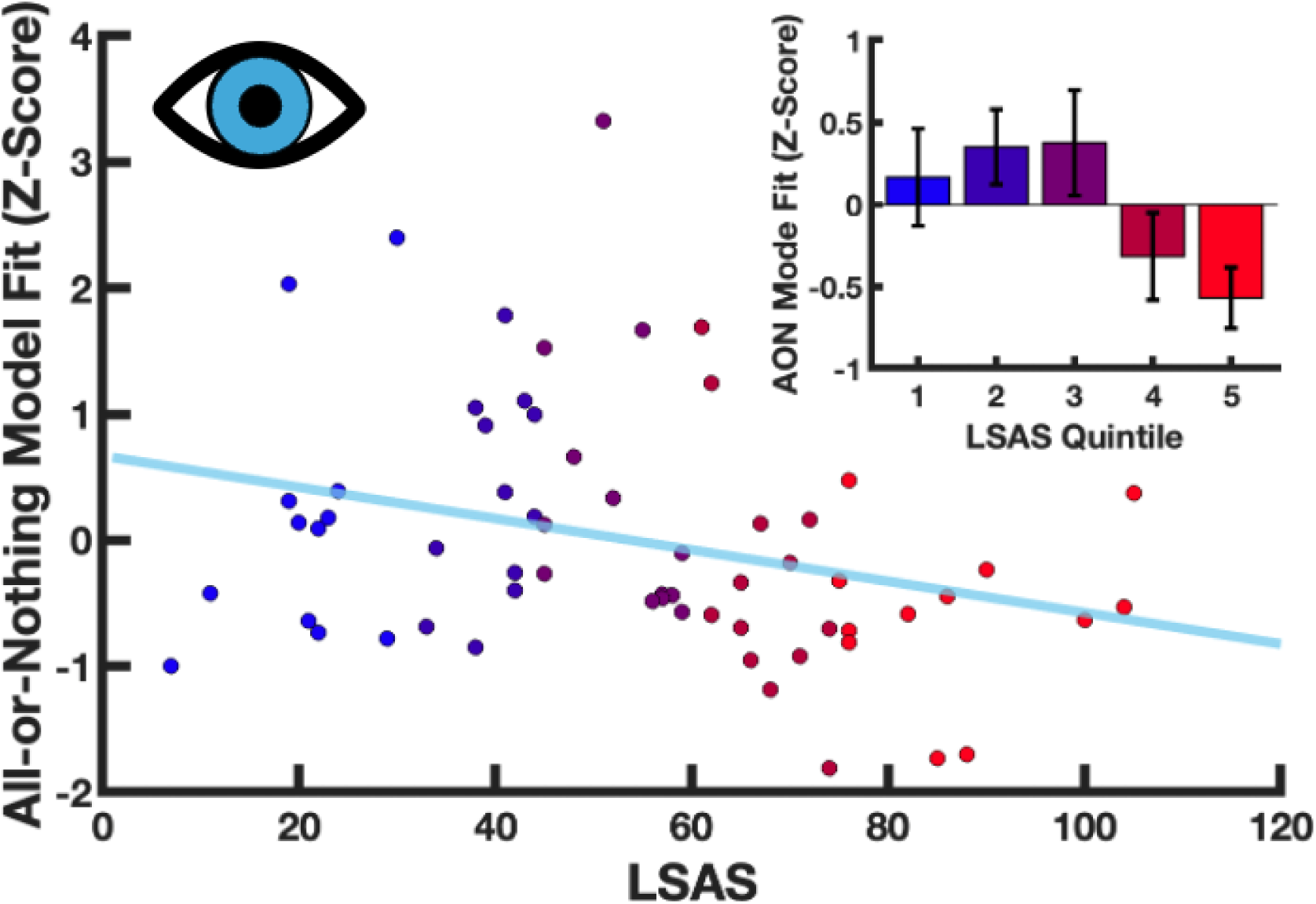
*Pupil Dilation Response Gradient Fit to All-or-Nothing Model against LSAS Ratings*. The inner product was taken between each participant’s condition-wise pupil dilation response pattern and the all-or-nothing pattern. The inner products were then Z-scored across participants, plotted against LSAS ratings, and color-coded based on LSAS quintile. Z-scored inner product averages and standard errors were calculated with respect to each LSAS quintile (*Top Right*).

Average self-reported valence and arousal ratings during early and sustained conditioning across the sample are shown in figure 12. Valence and arousal response gradients were expected to show increased generalization or discrimination as a function of LSAS ratings (M = 54.21, SD = 25.33, Min = 7, Max = 122). No significant relationship was found between social anxiety and any of the self-report measures (Table 1).

**Figure 12.**
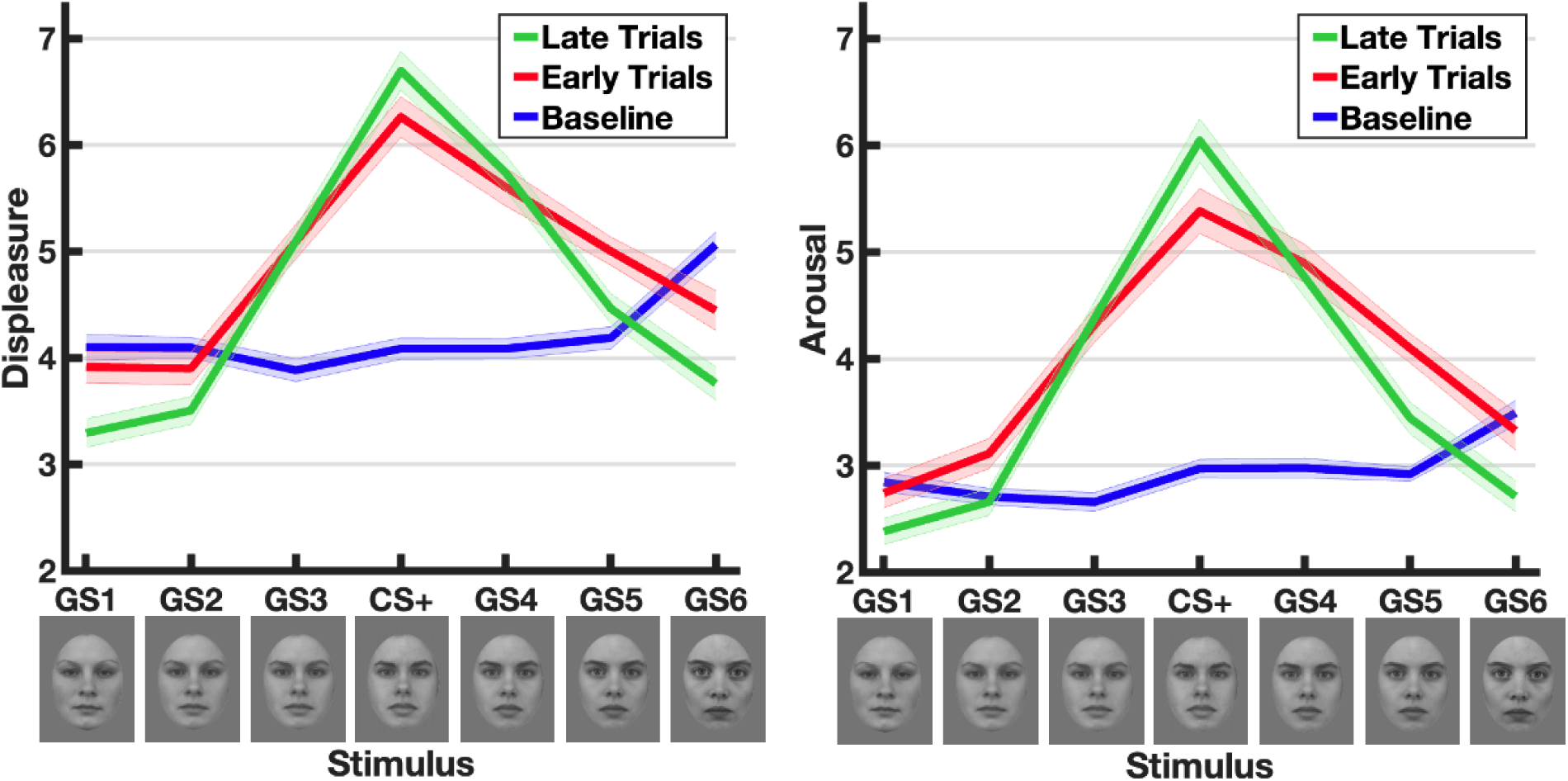
Average Valence and Arousal Ratings. Self-reported valence and arousal ratings were taken prior to conditioning (baseline), following an early trial (trial 87), and following a late trial (trial 168). Shaded error bars display within-subject variability (Cousineau, 2005).

**Table 1.** Rank-Correlations Between Valence and Arousal Inner Products and Social Anxiety Ratings. For each rating period during conditioning, inner products were computed between self-reported valence and arousal response patterns and both the generalization model and the all-or-nothing model. After Z-scoring across participants, each set of inner products was rank-correlated with LSAS ratings. No significant correlation was found for any rating period (all p’s < 0.025).

| Valence |  |  | Arousal |  |  |
| --- | --- | --- | --- | --- | --- |
| Rating Period | Generalization Model Fit | All-or-Nothing Model Fit | Rating Period | Generalization Model Fit | All-or-Nothing Model Fit |
| Early | 0.12 | 0.07 | Early | 0.15 | 0.16 |
| Late | 0.13 | 0.17 | Late | 0.08 | 0.01 |

## Discussion

SAD has been shown to lead to an increased sensitivity to faces, alongside a decreased ability to discriminate threatening and neutral faces (McTeague et al., 2018). The present study examined whether psychophysiological response patterns to a gradient of similar faces were altered during an aversive generalization paradigm as a function of self-reported social anxiety severity. Response gradients—and their relationship to social anxiety—varied between the different measures utilized in this experiment. This accords with the notion that human behavior is contingent on different underlying, non-linearly related systems, whose function varies by context, and based on the strategic and tactical needs of the organism (Lang, 1994). The present findings will be discussed separately for each measure in the following sections.

### ssVEP

In the present study, ssVEP amplitude was used to measure threat-related biases in visuocortical activation across stimuli varying in similarity to the CS+. Contrary to findings from Stegmann et al. (2020), ssVEP response patterns increasingly exhibited a generalization, rather than sharpening, pattern as social anxiety increased. One plausible explanation for this discrepancy pertains to the participant characteristics relative to their study. Stegmann and colleagues recruited entirely from the undergraduate student population; for this study, we took measures to additionally recruit from treatment-seeking members of the community. In support of these findings, McTeague et al. (2018) found a non-monotonic relationship between social anxiety and ssVEP amplitude differences between angry and neutral faces. This trend showed an increased response to angry faces relative to neutral as social anxiety increased, while the opposite pattern was found (greater response to neutral relative to angry faces) from participants with the most severe social anxiety. In exploratory quintile analyses, a similar pattern was found in the high LSAS quintiles, where visuocortical responses to one or more of the CS+-adjacent stimuli (GS3 and GS4) were similar to or greater than the response to the CS+. These quintile averages exhibit deviations from the sharpening pattern, which was more prominent in the low anxiety quintiles.

Modulation of visuocortical signaling as a function of motivational significance has been hypothesized to result from re-entrant feedback from frontal cortices (Petro et al., 2017). Successful treatment of social anxiety has been shown to increase prefrontal BOLD activation while altering visuocortical and limbic/paralimbic BOLD activation (Doehrmann et al., 2013; Goldin et al., 2013; Månsson et al., 2015). Additionally, recovery from SAD symptoms is predicted by greater activity in prefrontal and higher-order visual areas in response to threat (Doehrmann et al., 2013; Klumpp et al., 2013). In line with these findings, the present work suggests that visuocortical generalization patterns may be the result of altered or disrupted re-entrant feedback. Follow-up studies utilizing concurrent EEG-fMRI will be needed to clarify the relationship between visual cortices and frontal regions in the context of aversive generalization.

The results of the present study may also deviate from previous findings due to differences in experimental design. Most studies recording ssVEPs during aversive generalization preceded stimulus generalization with a differential conditioning phase, in which only the CS+ and a single CS– are presented (Stegmann et al., 2020; Stegmann & Gamer, 2025; Antov et al., 2020). However, other work has found that increased discrimination training prior to stimulus generalization attenuates generalization of self-report and autonomic responses (Ginat-Frolich et al., 2017). Additionally, Dunsmoor and LaBar (2013) found, in a study utilizing a symmetrical color gradient, that prior differential conditioning using the CS+ and a GS on one end of the gradient decreased autonomic and behavioral generalization across that half of the stimulus gradient. Based on these results, the present study forewent this initial differential conditioning phase to increase the degree of generalization for behavioral and autonomic measures. Doing so may also decrease the precision of CS+ identification by the visual system in persons with social anxiety, and magnify attentional resources dedicated to adjacent stimuli. Further work directly comparing the inclusion and exclusion of an initial differential conditioning phase could more conclusively elucidate these contrasting effects of social anxiety on generalization learning in visual cortex.

### Pupil Dilation

Against our hypotheses, results only showed a decreased fit of pupil dilation response gradients to the all-or-nothing model as social anxiety increased. Decreased relative fit to the all-or-nothing model may be interpreted as decreased discrimination of the CS+ by the pupil dilation response. Although the quintile-wise contrast results support a prominent linear trend, the distribution of all-or-nothing inner products against LSAS scores suggest a step function by which participants may be grouped into low social anxiety, high CS+ discriminators and high anxiety, low CS+ discriminators. In the high anxiety group, participants on average either exhibited broad generalization across the response gradient or did not respond preferentially to the CS+. Abundant work, primarily using cardiac measures, has found that persons with SAD show signs of autonomic dysregulation during rest and in response to stressors (Asbrand et al., 2022; Alvares et al., 2013; Schmitz et al., 2013). The present results support the use of task-evoked pupillary responses to measure autonomic dysregulation in response to stressors. In particular, increased pupil dilation responses to safe cues relative to threat suggest increased sympathetic reactivity in persons with SAD (Bradley et al., 2017; De Zorzi et al., 2021). It remains an open question whether this finding is exclusive for phobia-specific stimuli, or if these results would hold for all stimulus gradients, whether for persons with SAD or other anxiety disorders.

The expected and, prima facie, consistent finding that response gradients would also increasingly generalize as social anxiety increased, was not found. This may indicate a shortcoming of the model-fitting methods employed here and may be explained descriptively by the quintile-wise averages. While the all-or-nothing model only predicted maximal responses to the CS+, the generalization model predicted a specific Gaussian pattern of stimulus generalization. Quintile-wise averages indicate that although some high anxiety participants exhibited the latter pattern, the highest anxiety participants deviated from Gaussian generalization while exhibiting decreased stimulus specificity. This indicates that additional metrics of generalization learning may be preferable for characterizing stimulus generalization patterns when weight vectors are too stringent (Ahumada et al., 2025; Stegmann et al., 2024).

### Ratings

While average valence and arousal ratings displayed generalization across the stimulus, these response patterns – contrary to our hypotheses – were not significantly altered as a function of social anxiety. These results resemble those of previous studies, which have found either a small effect or non-significant of social anxiety on generalization of behavioral ratings (Ahrens et al., 2016; Stegmann et al., 2021; Reutter et al., 2025).

Altogether, these findings demonstrate the importance of characterizing psychopathology with psychophysiological recordings, rather than exclusively relying on self-report measures (Lang & McTeague, 2009). Incorporating the former into psychiatric assessment would decrease the burden on psychiatric patients, who may vary in their abilities to produce descriptions of their symptoms which conform with prevailing clinical nosologies (Morris et al., 2022). In the context of SAD, discrepancies in generalization of conditioned fear relative to healthy controls may not be apparent in the self-report ratings, despite clear differences in physiological behaviors. For this reason, it may be beneficial to implement aversive generalization paradigms utilizing psychophysiological recordings to inform diagnosis and treatment for SAD.

## Conclusion

The present work found that social anxiety moderately increases fit of visuocortical responses to a generalization model and decreases CS+ discrimination of pupil dilation responses. Overall, these results suggest that, in patients with SAD, both perceptual and autonomic systems increasingly attribute threat-related information to neutral stimuli in virtue of their similarity. This work may help increase understanding of the role of perception and learning on the etiology and treatment of SAD. Further work incorporating concurrent fMRI could explain how these differences in response gradients relate to differences in functional connectivity, allowing for a greater understanding of how these processes are dysregulated in SAD. Additionally, work using a larger sample size and greater variability in clinical symptomology could elucidate interactions between different psychopathologies on aversive generalization in SAD patients.

