## Supplemental Figures for "Social Anxiety Increases Autonomic and Visuocortical Generalization of Conditioned Aversive Responses to Faces"

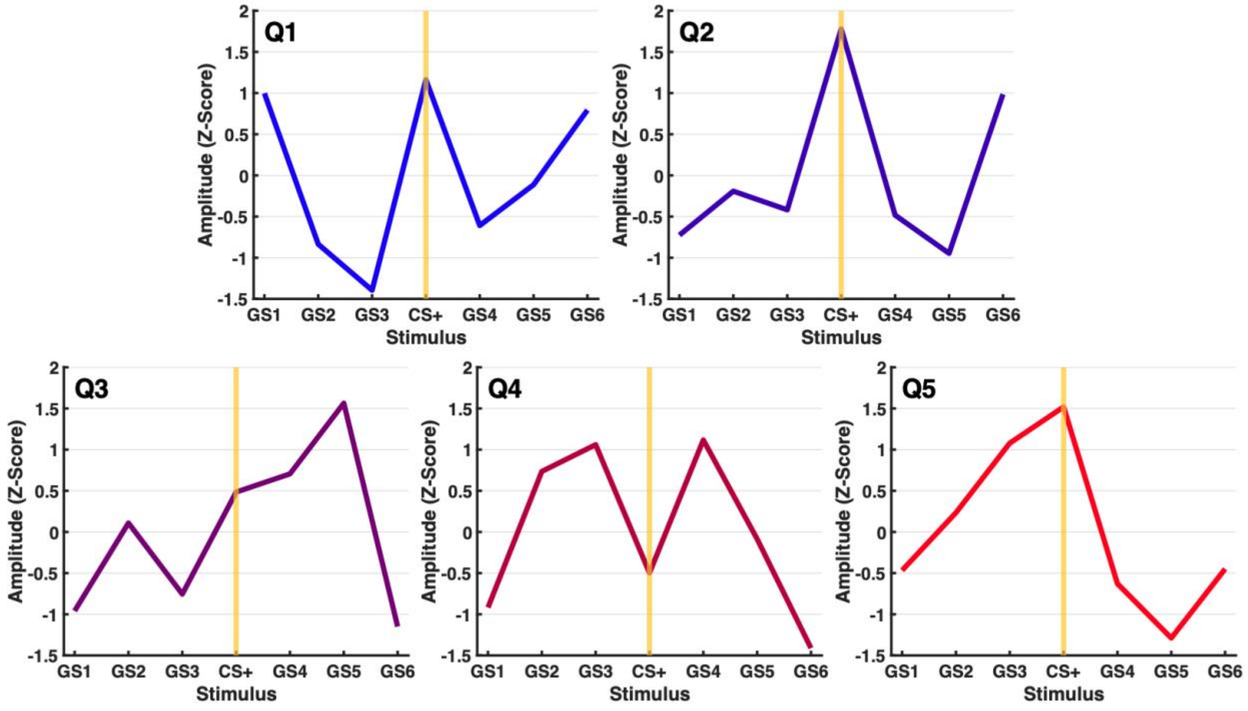


*Supplementary Figure 1*. *ssVEP Response Gradients Grouped by LSAS Quintile*. ssVEP data were averaged within quintiles determined by LSAS ratings. Each quintile average response gradient was Z-scored to enable comparison between groups.

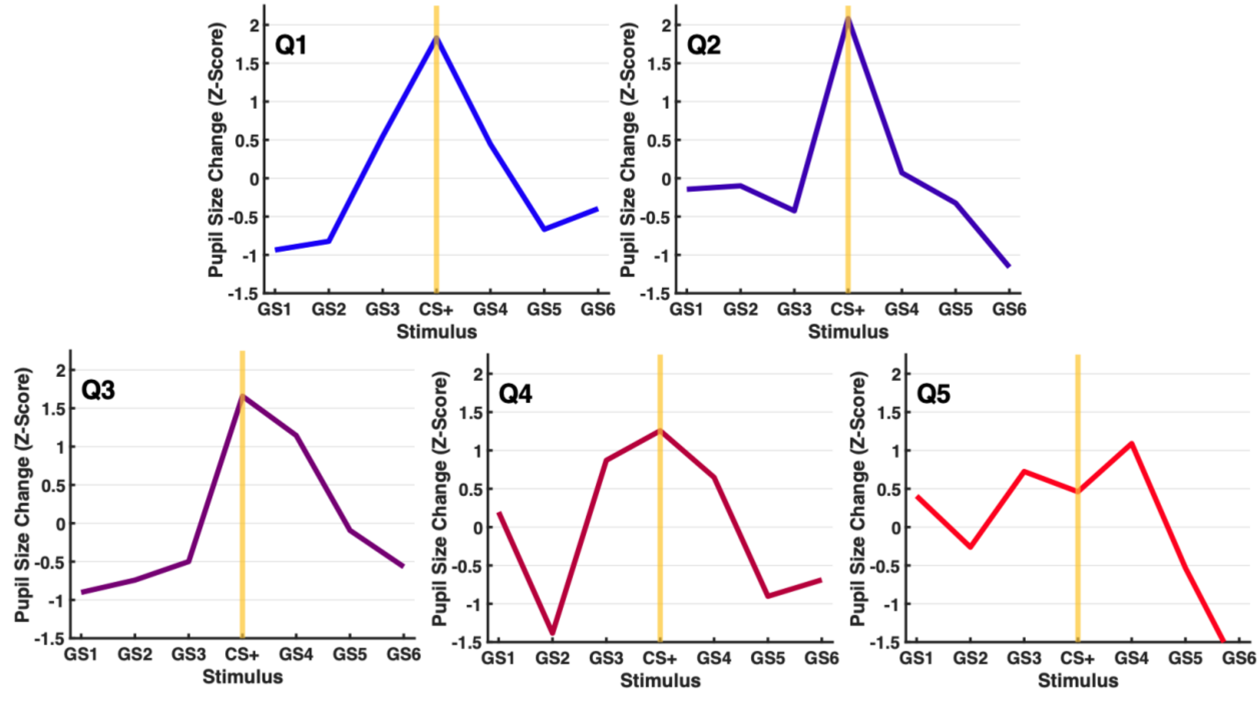


*Supplementary Figure 2*. *Pupil Dilation Response Gradients Grouped by LSAS Quintile*. Pupil data were averaged within quintiles determined by LSAS ratings. Each quintile average response gradient was Z-scored to enable comparison between groups.
